# Yolk steroid concentrations decline in Japanese Quail (*Coturnix japonica*) eggs throughout incubation

**DOI:** 10.1101/2025.06.02.657472

**Authors:** S.K. Winnicki, M.E. Hauber, R.T. Paitz, T.J. Benson

## Abstract

The development of avian embryos is dependent not only on their genetic background but also on both external conditions and the maternal resources deposited into eggs. Specifically, maternally derived steroids in egg yolks have been shown to influence morphological development in avian offspring, but the effect of most yolk steroid hormones and their metabolites on embryonic growth and development remain poorly explored. We tested relationships between eggshell maculation, surface temperature, hormone concentrations before and during incubation, and embryonic growth in Japanese Quail (*Coturnix japonica*) eggs developing while artificially incubated in duplicate incubators. We detected 10 steroid hormones in the yolks during development and concentrations of all hormones declined to Day 15 (∼2 days before hatch). Steroid declines at Day 9 of incubation were related to eggshell surface temperature, with warmer eggs having higher concentrations of androgens and progestogens, and incubator assignment, as eggs were warmer on average in one incubator than the other. However, eggshell surface temperature was not related to eggshell maculation, embryonic size, or embryonic heart rate during development. These results provide evidence of yolk steroid concentration declines through incubation and suggest that environmental conditions (such as temperature or light) could alter embryonic hormone uptake and/or metabolism.

## Introduction

Developing organisms grow at different rates both within and among species (Starck and Ricklefs 1998) with consequences for the organisms’ survival and fitness (Monaghan and Ozanne 2008). Birds complete most of their development outside of the maternal body and exhibit a wide range of developmental modes both prior to and following hatching from the eggshell (Starck and Ricklefs 1998); for example, birds grow faster than any other extant vertebrates, and their structure-specific growth patterns vary among species, populations, and individuals (McCarty 2001). Although many of the observed differences are likely genetic and epigenetic in origin, diversity in growth rates may also result from variation in early environmental conditions, including maternal programming through the manipulation of the egg’s content (West-Eberhard 1989).

The maternal hormones deposited into bird eggs have been of particular interest to researchers investigating ways that parents might tune offspring growth to meet environmental challenges (Groothuis et al. 2019). Initial concentrations of egg-yolk hormones, for example, can impact post-hatch phenotype; accordingly, natural or injected androgens in canary (*Serinus canaria domestica*) eggs enhanced post-hatch growth and influenced the nestlings’ begging behavior (Schwabl 1996). However, these patterns are not always consistent across other bird species and studies (Groothuis et al. 2019).

Some previous work on avian maternal hormone investment has focused on a subset of hormones including thyroid hormones (Ruuskanen and Hsu 2018), androgens (Eising et al. 2001, Eising and Groothuis 2003, Groothuis et al. 2019), and glucocorticoids (Bowers et al. 2019), despite the dozens of other hormones and their metabolites present in freshly laid bird eggs, including their yolks (Merrill et al. 2019). Critically, increasing evidence suggests that maternally invested hormones are rapidly metabolized by embryos at the onset of development (von Engelhardt et al. 2009, Paitz et al. 2010, Paitz et al. 2020), yet the majority of these metabolites, and their impacts (if any) on embryonic growth, remain poorly characterized in the context of maternal effects (but see Kumar et al. 2018, Wang et al. 2023). For this reason, it is critical that time-series studies examine both the ontogenetic yolk hormone concentration shifts and co-occurring embryonic developmental metrics in the incubated avian egg.

Developing avian embryos are also subjected to variation in the external environment during development. While the eggshell acts to separate and protect the embryonic environment from the external environment, it also allows for the transfer of gases, water vapor, some microbes, heat energy, and some light (Roberts and Brackpool 1994, Maurer et al. 2014, Lahti and Ardia 2016). For example, lighted incubation shortens incubation time in turkeys (*Meleagris gallopavo f. domestica*, Fairchild and Christensen 2000), and broiler chicken (*Gallus gallus domesticus*) embryos incubated under 24-hour green light gained more weight than birds incubated in the dark (Rozenboim et al. 2013). Photoperiod may interact with steroid hormones to influence hatchling physiology (Deviche et al. 2001), including hatchling blood androgen levels (Yu et al. 2018) and corticosterone levels (Özkan et al 2012). If light exposure should alter embryonic development by altering embryonic uptake and metabolism of maternally-derived yolk steroids, this effect could be further impacted by both eggshell characteristics and egg temperature. Pigmentation in eggshells can absorb harmful UV-B radiation while reducing the reflectance of the eggshell, thereby warming it (Magige et al. 2008). Because eggs incubated at temperatures just 1°C lower than conspecifics produce slower-growing embryos (Lourens et al. 2005) and eggs with greater maculation could absorb more thermal energy from light, embryo development may always be faster in light-exposed eggs with greater maculation. However, eggshell pigmentation may be correlated with egg contents (Navarro et al. 2011), including yolk hormones (López-Rull et al. 2008), and any temperature variation could impact embryonic hormone production and metabolism because associated enzymes are temperature-dependent (Yalcin et al. 2022). Disentangling the relationships between eggshell pigment maculation, eggshell surface temperatures, maternally-derived egg hormones, hormone metabolism/uptake, and embryonic development during lighted incubation requires assessing temperature, maculation, and hormones across development in eggs experiencing.

We assessed yolk steroid hormone concentrations in unincubated eggs and progressively incubated eggs and the developing embryos of Japanese Quail (*Coturnix japonica*) in duplicate incubators and related these metrics to variation in individual eggshell maculation, eggshell surface temperatures, and incubator-level light exposure to the growth and development (heart rate) of the embryos and the concentrations of yolk steroid hormones at multiple points in development. Since domestic chicks (*Gallus domesticus*) developing under light gain more mass than chick developing in the dark (Rozenboim et al. 2013), we predicted that quail embryos developing in more light would be (1) larger overall by the end of the developmental period and (2) lead to embryos taking up maternally deposited steroid hormones during development more rapidly than in the half-dark treatment. If eggshell maculation is an indicator of egg contents, we predicted (3) that eggshell maculation would be related to the concentrations of unincubated yolk steroids. If eggshell maculation impacts surface temperature to influence embryonic development, we also predicted (4) that eggs with greater maculation would have higher surface temperatures throughout development and that the resulting embryos would reach larger sizes, especially in eggs exposed to constant light. Finally, if eggshell surface temperatures alter embryonic hormone uptake and/or metabolism, we predicted (5) that eggs with warmer surface temperatures would also have accelerated yolk steroid declines over development and larger embryos relative to eggs with lower surface temperatures.

## Methods

We obtained fertilized Texas A&M line Japanese Quail eggs from a commercial breeding farm (Purely Poultry, Fremont WI, USA) and measured the length and width of each egg (to the nearest 0.1 mm using calipers) and fresh mass of the incubated egg (to the 0.01 g), took standardized photos of the eggs for eggshell maculation analyses (Canon Powershot Sx60, Canon Inc, Huntington NY, USA), and froze N=6 eggs at −20°C before incubation to measure initial hormone content and concentrations. Because of the presence of infertile eggs, the subsequent sample sizes for each analysis vary throughout the experiment; a visualization of sample sizes can be found in Figure 1.

**Figure 1.**
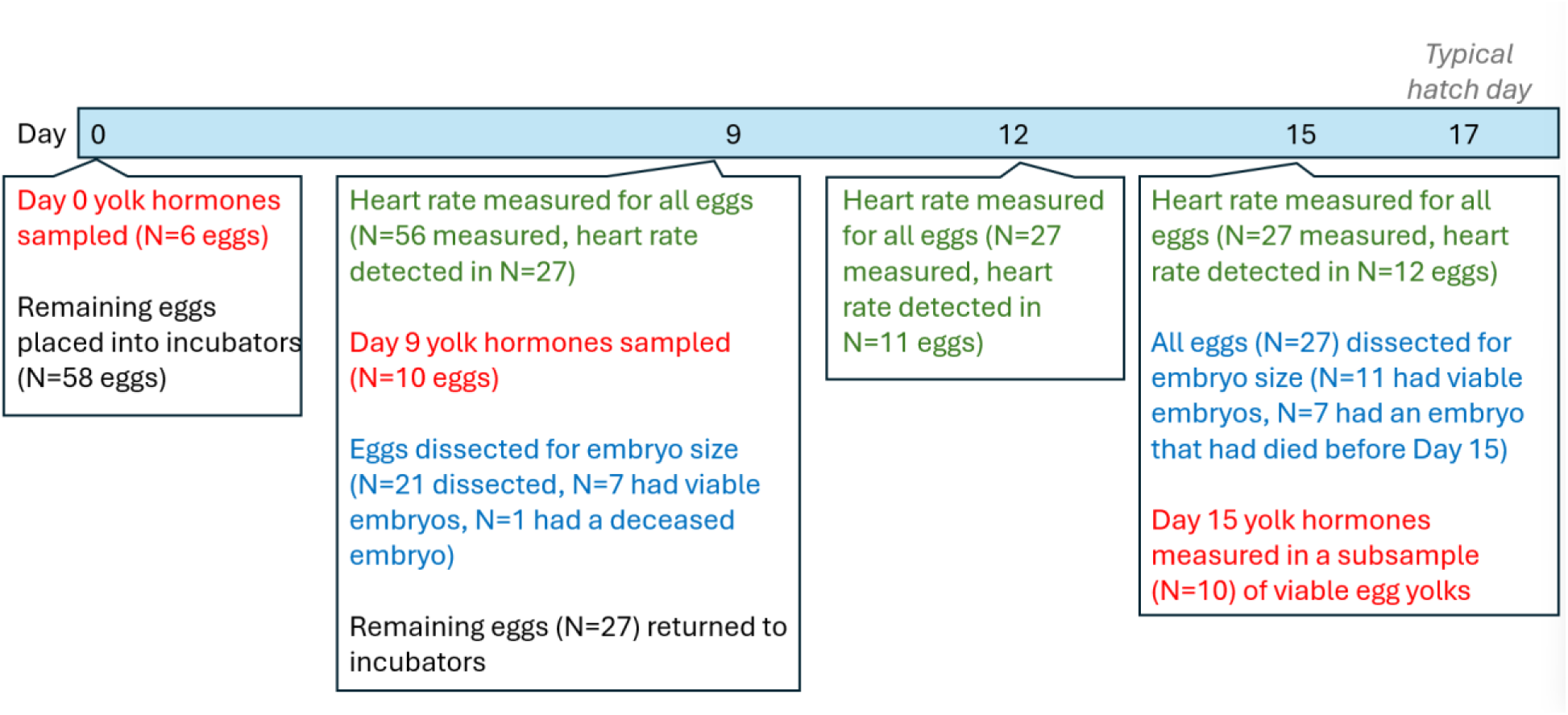
Quail egg-sampling design, indicating the days of incubation when sampling occurred and the type of sampling (freezing eggs for yolk steroid analysis, dissecting eggs for embryonic size measurements, and using heart monitors to measure embryonic heart rate in living eggs).

The clutch of origin and the laying order of these eggs were unknown, so we gave eggs an individual label with a Sharpie marker and randomly assigned eggs to one of two clear-topped incubators (Rcom Pro 20, Rcom Incubators USA, Wichita KS, USA) set at 37.5°C and 45% humidity and to turn the eggs 60° every 40 minutes. Both incubators were placed under a wooden box to control light; the duplicate incubators were lit with natural-light 6500k LEDs (Minger, Govee Moments, Hong Kong, HK) at ∼545 lux, consistent with the amount of light that would naturally hit a forested ground-nest (Shirley 1929, Janick 1972) but below extreme light intensities used in previous studies of development (e.g. 1340-1730 lux, Shafey and Al-Mohsen 2002). Eggs in one incubator were exposed to 24 hours of light while eggs in the other incubator were exposed to 12 hours on/off, controlled by an automatic timer. We acknowledge the limitations of our study by having used different light treatments in each of the two incubators; we therefore model effects of incubator identity rather than light treatment, as other factors that may have differed between the two incubators (such as temperature, see Results below) may have also impacted embryonic development.

On Days 9, 12, and 15 of the eggs’ ∼17 day incubation period we measured the heart rates of the embryos in the remaining eggs using noninvasive ballistocardiogram monitors (EggBuddy, Avitronics, Cornwall, UK) modified to record the data in real-time over a 5-minute window (Di Giovanni et al. 2023). Prior to placing the eggs into the EggBuddy we removed the eggs from the incubator one at a time, immediately measuring their eggshell surface temperature using a handheld infrared visual temperature monitor (FLIR TG165, Teledyne FLIR LLC, Wilsonville, OR, U.S.A.). When we modeled the effect of temperature on development (below) we used the average of all temperature measurements we took for an individual egg. Since some eggs were sacrificed after measurements on Day 9 the temperature data for these individuals was the Day 9 eggshell surface temperature alone, but for eggs sacrificed on Day 15 the temperature data used was the average of eggshell surface temperatures sampled on Days 9, 12, and 15. Egg manipulations always occurred during periods when the lights were on. We then placed each egg into the EggBuddy and recorded five minutes of activity before returning the eggs to the incubators. In the R statistical software (R Core Team 2013) we calculated the average heart rate measured over those 5 minutes.

On the 9^th^ day of incubation, we froze 10 eggs (5 from each incubator) for hormone analyses. We dissected an additional 21 unfrozen eggs (10-11 from each incubator) to measure both the fresh eggshell thickness and the embryo size. Since some of the eggs sampled did not contain a living embryo, we measured 7 viable embryos total (N=3 from the 24 hour on incubator, N=4 for the 12 hours on/off incubator). We measured the embryo’s wing, tarsus length, and eye diameter (±0.1 mm) with calipers when embryos were present. We also recorded each embryo’s mass (±0.01 g). Using a linear gauge we quantified the thickness of each dissected embryo’s eggshell to the nearest 0.001 mm on the top, bottom, and side of the shells, repeating the measurements three times at each location for each egg. We did not sex the quail embryos in this study, as our sample sizes would have been too small per age category for meaningful statistical analyses.

On the 15^th^ day of incubation (out of a total incubation period of 17 days in Japanese quail) we froze the rest of the eggs (27 eggs total, N=13 for 24 hour incubator, N=14 for 12 on/off incubator). Seven of these eggs contained embryos that had begun development but had died prior to Day 15 and 10 eggs contained no embryos, leaving us with 11 living embryos (6 from the full light incubator and 5 from the 12 hour on/off incubator). Because we did not have enough viable eggs to sample 5/incubator for hormones and measure 5/incubator unfrozen eggs for embryos and eggshell thickness, as we had done on Day 9, we instead froze all the eggs. We measured the embryos in all of the frozen eggs and randomly chose 5 frozen yolks from each of the two incubators for the costly hormone analyses. We did not measure eggshell thickness of Day 15 eggs because the freezing process crumbled the eggshells.

We prepared yolk samples and extracted steroids using methods of Merrill et al. (2019), aiming to quantify the quantity and concentrations of 29 yolk steroid hormones, including the *estrogens* estrone and estradiol; *androgens* androstenedione, testosterone, dehydroepiandrosterone (DHEA), etiocholanolone, and 11-ketotestosterone; *glucocorticoids* 11-dexoycortisol, cortisone, 5β-dihydrocortisone, 20β-dihydrocortisone, cortisol, 5β-dihydrocortisol, 5β-tetrahydrocortisone, 5β-tetrahydrocortisol, β-cortol, corticosterone, 5β-tetrahydrocorticosterone, 11-dehydro-tetrahydrocorticosterone, 5β-corticosterone, deoxycorticosterone, 11α-hydroxyprogesterone, and 20β-dihydrocorticosterone; and *progestogens* progesterone, pregnenolone, 17α-hydroxypregnenolone, dihydroprogesterone (DHP), pregnanedione, and pregnanolone.

We extracted yolks from the frozen eggs, homogenized them, added 0.5 g of yolk to 4 mL of 100% methanol, stored them at −20°C overnight, and centrifuged them at 2000 r/min at 4°C for 15 minutes. We extracted the supernatant, added 46 mL water to dilute the samples further steroid extraction. We performed solid-phase extraction on the supernatant using C18 Sep-pak cartridges (Waters Ltd., Watford, UK) by charging the cartridges with 5mL 100% methanol, rinsing them with water, loading the sample, running the loaded cartridges under vacuum pressure (3-5 Hg), rinsing again with water, and eluting the samples with 5mL ether. Samples were dried overnight and submitted to the Roy J. Carver Biotechnology Center (University of Illinois at Urbana-Champaign) Metabolomics Laboratory for LC-MS-MS analysis on a 5500 QTRAP LC-MS-MS system (AB Sciex, Foster City, California, USA), which quantifies all 29 hormones in a single run if the hormones are above the detection threshold of ∼10pg (Merrill et al. 2019, Hauber et al. 2020). Recoveries ranged from 56% for 17OH-hydroxypregnenolone to 82% for pregnenedione. We analyzed all the hormone results as the measured hormone concentrations (ng/g of yolk).

Using ImageJ (Schneider et al. 2012) we converted the photo of each egg taken along the long axis lying flat to a light/dark binary image and quantified the proportion of the visible eggshell area covered by dark maculation (spotting).

We used linear models in program R (R Core Team 2013) to compare the maculation percentage, length, width, mass, and individual hormone concentrations of eggs at Day 0 assigned to the two incubators. We used chi-squared analyses in R to assess whether there was an incubator effect in the eventual fate of eggs (viable or inviable at sampling point Day 9 and Day 15). We averaged all shell-temperature measurements for each individual egg over the course of the experiment (measured on day 9 of incubation or days 9, 12, and 15 of incubation) and used linear models to assess the effect of incubator and eggshell maculation percentage on average eggshell temperature. We assessed the relationship between eggshell maculation and maternally-deposited hormones by running univariate models of the effect of eggshell maculation percentage on the concentration of each steroid hormone detected on Day 0 (pre-incubation).

To assess the effect of sampling day, eggshell temperature, incubator assignment, and eggshell maculation on the concentrations of all hormones detected in the egg yolks across development, we used multivariate analysis of variance (MANOVAs) with scaled hormone measurements on sampling days 0, 9, and 15 as dependent variables. Due to sample size limitations we could not include all the predictors in one model, so we instead constructed five MANOVAs of hormone concentration: one MANOVA with the sampling day; one MANOVA with additive effects of sampling day and incubator assignment and an interaction between sampling day × incubator assignment; one MANOVA with the sampling day and an additive effect of average eggshell surface temperature; one MANOVA with sampling day and an additive effect of eggshell maculation; and a final MANOVA with an effect of sampling day, the main effects of eggshell maculation and incubator assignment, plus the interaction between eggshell maculation and incubator assignment. We present effect sizes as eta-squared values with 95% confidence intervals. We used a separate set of MANOVAs, with the same predictor structure, to investigate the impact of sampling day, incubator assignment, eggshell maculation, and eggshell surface temperature on embryo size measurements on Day 9 and Day 15 of incubation, using the embryo tarsus, bill, wing, eye, and mass measurements as the dependent variables. When these MANOVA results were significant at α=0.05 we performed a linear discriminant analysis with the same predictors as the MANOVA to explore the response variables that varied most between the treatment groups. LDAs were performed with the R package *MASS* (Venables and Ripley 2002).

Because we have yolk hormone concentrations and embryo size measurements for the same eggs on Day 15, we were able to relate the suite of hormones present in the individual egg to the size of the embryo in that same egg, unlike the eggs sampled on Day 9. Since both hormone concentrations and embryo size are multivariate datasets, we reduced both down to individual principal components by performing separate principal components analyses on yolk hormones and embryo size, using the R package *FactoMineR* (Lê et al. 2008). We then investigated an effect of the resulting yolk hormone principal components on the embryo size principal components using linear models.

## Results

The eggs assigned to each incubator did not vary in maculation (mean±SE= 137.77 ± 7.89, *F*_1,55_=2.30, *P*=0.135), egg length (mean±SE= 35.73 mm ± 0.20, *F*_1,66_=0.05, *P*=0.825), width (mean±SE= 27.33 mm ± 0.11, *F*_1,66_=1.04, *P*=0.311), or initial mass (mean±SE= 14.64 g ± 0.17, *F*_1,66_=0.16, *P*=0.687).

At Day 9 of incubation 61.3% of embryos had survived and fate was not dependent on the incubator in which the eggs were held (χ^2^_1_=0.93, *P*=0.335). Eggshell thickness in eggs with viable embryos at Day 9 did not differ between the incubators (eggshell top: mean±SE= 0.085 mm ± 0.002, *F*_1,6_=0.243, *P*=0.640; middle: mean±SE= 0.081 mm ± 0.002, *F*_1,6_=1.205, *P*=0.314; bottom: mean±SE= 0.079 mm ± 0.002, *F*_1,6_=0.503, *P=*0.505). At Day 15, 40.7% of the embryos had survived and embryo fate was not dependent on incubator (χ ^2^_1_=0.03, *P*=0.873). Average eggshell surface temperatures (across Days 9, 12, and 15 of incubation) were ∼1 degree Celsius higher in the 24-hr on incubator than in the 12-hour on/off incubator (β±SE: 1.03±0.45, *F*_1,55_=-5.18, *P*=0.027). Eggshell maculation did not impact the average surface temperature of each egg (β±SE: 0.002±0.004, *F*_1,55_=0.30, *P*=0.586).

### Steroid Hormone Ontogenetic Shifts

We detected 9 of the 29 targeted hormones in the unincubated (Day 0) egg yolks: androstenedione, testosterone, dehydroepiandrosterone (DHEA), etiocholanolone, progesterone, pregnenolone, 17α-hydroxypregnenolone, pregnanedione, and pregnanolone. Day 0 egg hormone concentrations did not vary between eggs randomly assigned to the incubators, nor did they vary with eggshell maculation (Supplementary Table S1).

All 9 hormones were still present at detectable quantities in the yolks at Day 9 of incubation and we additionally detected 5β-tetrahydrocortisol in four Day 9 eggs. By Day 15 of development, only androstenedione, progesterone, pregnenolone, pregnanedione, and pregnanolone were detectable in the egg yolks (Table 1). Yolk hormone concentrations decreased over development (η^2^[95%CI]: 0.99[0.97, 1.00], *F*_10,13_=86.93, *P*<0.001, Figure 2) but there was no significant additive effect eggshell maculation (η^2^[95%CI]: 0.26[0.00, 1.00], *F*_10,14_=0.48, *P*=0.875), or eggshell maculation × incubator assignment (η^2^[95%CI]: 0.59[0.00, 1.00], *F*_10,12_=1.74, *P*=0.181).

**Figure 2.**
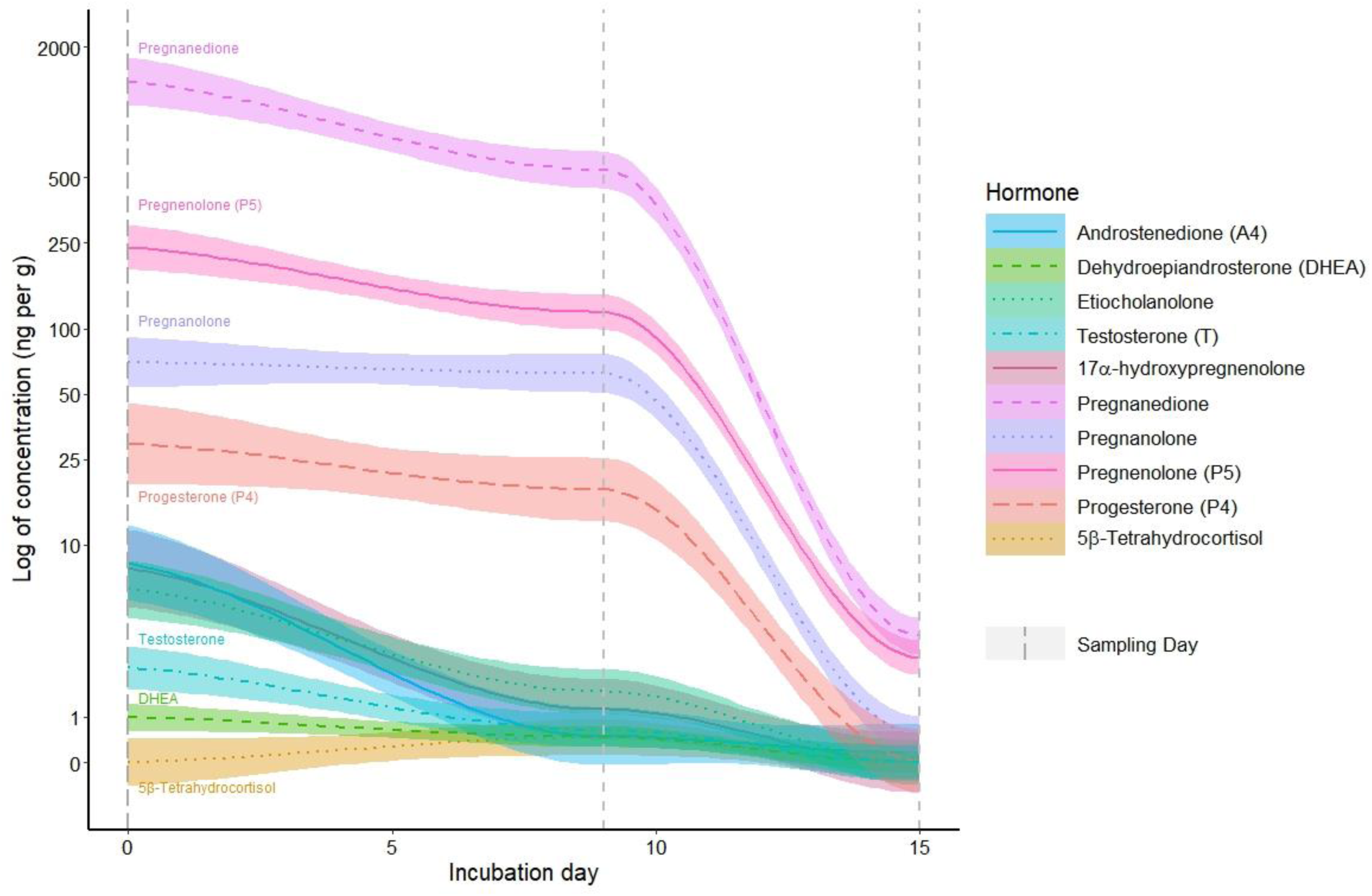
Yolk hormone concentrations (ng/g) detected on sampling days 0, 9, and 15 of incubation across all sampled quail eggs. The lines represent the predicted average concentration of each hormone and the shaded areas represent 95% confidence intervals, while the dotted vertical lines represent sampling days.

**Table 1.** Range, mean, and standard deviation of quail yolk hormone concentrations (in ng/g) in eggs sampled on days 0, 9, and 15 of incubation in the two incubators (exposed to 24 hours of light or 12 hours on/off). There was no significant effect of incubator assignment on hormone declines (F_10,14_=0.93, *P*=0.535).

|  | Day 0 | Day 9 |  | Day 15 |  |
| --- | --- | --- | --- | --- | --- |
|  | no incubation | 12 on/off incubator | 24 on incubator | 12 on/off incubator | 24 on incubator |
|  | N=6 | N=5 | N=5 | N=5 | N=5 |
| **Androstenedione (A4)** |  |  |  |  |  |
| range | 4.84–15.76 | 0 | 0–4.82 | 0 | 0–2.42 |
| mean ± SD | 8.89 ± 4.07 | 0.00 ± 0.00 | 1.55 ± 2.23 | 0.00 ± 0.00 | 0.48 ± 1.08 |
| **Dehydroepiandrosterone (DHEA)** |  |  |  |  |  |
| range | 0.29–2.16 | 0.22–1.01 | 0–0.58 | 0 | 0 |
| mean ± SD | 1.03 ± 0.72 | 0.71 ± 0.31 | 0.40 ± 0.25 | 0.00 ± 0.00 | 0.00 ± 0.00 |
| **Etiocholanolone** |  |  |  |  |  |
| range | 2.60–16.82 | 0.80–1.61 | 1.42–2.88 | 0 | 0 |
| mean ± SD | 7.62 ± 5.69 | 1.26 ± 0.34 | 2.12 ± 0.52 | 0.00 ± 0.00 | 0.00 ± 0.00 |
| **Testosterone (T)** |  |  |  |  |  |
| range | 1.04–4.44 | 0.24–0.85 | 0.16–2.78 | 0 | 0 |
| mean ± SD | 2.53 ± 1.30 | 0.47 ± 0.26 | 1.03 ± 1.08 | 0.00 ± 0.00 | 0.00 ± 0.00 |
| **17α-hydroxypregnenolone** |  |  |  |  |  |
| range | 4.34–10.40 | 0–4.24 | 0–4.66 | 0 | 0 |
| mean ± SD | 8.10 ± 2.55 | 0.85 ± 1.90 | 2.30 ± 2.22 | 0.00 ± 0.00 | 0.00 ± 0.00 |
| **Pregnanedione** |  |  |  |  |  |
| range | 828–1748 | 296–660 | 474–976 | 2.78–6.10 | 2.80–4.56 |
| mean ± SD | 1,434.33 ± 362.96 | 456.00 ± 137.69 | 690.80 ± 193.83 | 3.81 ± 1.38 | 3.53 ± 0.82 |
| **Pregnanolone** |  |  |  |  |  |
| range | 42.4–130.8 | 30.0–88.4 | 44.8–124.4 | 0.44–0.84 | 0.29–1.14 |
| mean ± SD | 74.97 ± 31.93 | 63.44 ± 23.23 | 69.92 ± 31.28 | 0.59 ± 0.17 | 0.57 ± 0.33 |
| **Pregnenolone (P5)** |  |  |  |  |  |
| range | 144.2–326.0 | 98.4–120.4 | 97.8–183.0 | 0.94–3.30 | 2.70–5.86 |
| mean ± SD | 245.37 ± 60.37 | 110.88 ± 10.19 | 132.24 ± 31.15 | 2.07 ± 0.85 | 3.70 ± 1.26 |
| **Progesterone (P4)** |  |  |  |  |  |
| range | 10.86–88.40 | 7.64–31.20 | 8.48–38.20 | 0–0.2 | 0–0.06 |
| mean ± SD | 36.18 ± 27.12 | 15.46 ± 9.43 | 27.68 ± 13.58 | 0.06 ± 0.09 | 0.02 ± 0.03 |
| **5β-Tetrahydrocortisol** |  |  |  |  |  |
| range | 0 | 0–2.94 | 0–2.74 | 0 | 0 |
| mean ± SD | 0.00 ± 0.00 | 0.68 ± 1.28 | 0.61 ± 1.20 | 0.00 ± 0.00 | 0.00 ± 0.00 |

However, in addition to sampling day, there was a significant additive effect of average eggshell surface temperature (η^2^[95%CI]: 0.82[0.07, 1.00], *F*_10,8_=3.55, *P*=0.042). A linear discriminant analysis of the effect of average eggshell surface temperature and sampling day produced only one linear discriminant score with trace values >10% (Supplementary Table S2). The first LD score described 99.9% of the variation in the hormone data (Supplementary Table S2) and split the samples by the sampling day (Supplementary Figure S1). While LD2 split the Day 9 samples by surface temperature (Supplementary Figure S1), this effect only explained 0.03% of the variation in the hormone data and was mostly driven by three eggs on Day 9 with higher surface temperatures than the average surface temperature measured across the eggs in the study (>31.3°C). The eggs with the higher average eggshell surface temperatures (> 31.3°C, N=3) had greater average concentrations of androstenedione, testosterone, and progesterone and lower average concentrations of 17α-hydroxypregnenolone and 5β-Tetrahydrocortisol than the eggs with the smaller average eggshell surface temperatures (< 31. 3°C, N=7), but the ranges of concentrations for each hormone overlapped substantially (Supplementary Table S3).

There was also a significant effect of the interaction between incubator assignment × sampling day on yolk hormone content. (η^2^[95%CI]: 0.75[0.26, 1.00], *F*_10,13_=3.86, *P*=0.013). The linear discriminant analysis of the effect of incubator × sampling day (Supplementary Table S4) showed similar results to the surface temperature hormone concentration LDA, showing the most difference between the hormone content of egg yolks on Day 9 (Supplementary Figure S2). Because the eggs with the highest eggshell surface temperatures (> 31.3°C) on Day 9 were in the same incubator (exposed to 24 hours of light) this incubator effect mirrors the previously described relationship between eggshell surface temperature and egg hormones.

### Embryonic Development

Tarsus, bill, wing, eye, and mass measurements increased over day measured (η^2^[95%CI]: 0.98[0.95, 1.00], *F*_5,12_=105.11, *P*<0.001) but that pattern was not influenced by an additional additive effect of incubator assignment (η^2^[95%CI]: 0.39[0.00, 1.00], *F*_5,10_=0.47, *P*=0.349), an additional interaction between incubator assignment × day measured (η^2^[95%CI]: 0.19[0.00, 1.00], *F*_5,10_=0.47, *P*=0.793) nor the additive effect of average eggshell surface temperature (η^2^[95%CI]: 0.12[0.00, 1.00], *F*_5,11_=0.29, *P*=0.909). While there was no significant effect of an interaction between eggshell maculation × incubator assignment (η^2^[95%CI]: 0.52[0.00, 1.00], *F*_5,9_=1.93, *P*=0.185), eggshell maculation did have a significant effect on embryo size as an additive fixed effect (η^2^[95%CI]: 0.64[0.10, 1.00], *F*_5,11_=3.99, *P*=0.026). A linear discriminant analysis of the effect of eggshell maculation and sampling day on embryonic size produced only one linear discriminant score with trace values >10% (Supplementary Table S5). The first LD score described 93.1% of the variation in the embryonic size data (Supplementary Table S5) and split the samples by the sampling day (Supplementary Figure S3). While LD2 split the Day 15 samples by maculation quartiles (Supplementary Figure S3), this pattern only explained 3.5% of the variation in embryonic size data and was mostly driven by two embryos on Day 15 in highly maculated eggs that had larger bill lengths (mean±SD= 7.35±0.64 mm, compared to 6.42±0.53 mm in eggs with less maculation) and smaller wing lengths (mean±SD =21.55±1.20 mm, compared to 24.01±2.12 mm in eggs with less maculation) compared with embryos growing in eggs that fell in the first three maculation quartiles.

Average heart rate increased with age (days since incubation start, β±SE: 16.27±3.11, *t*_21_=5.24, *P*<0.001, Supplementary Figure S4). Average heart rate was not significantly impacted by incubator (β±SE: 2.81±16.44, *t*_27_=0.17, *P*=0.866), average eggshell surface temperature (β±SE: −1.61±5.01, *t*_27_=-0.32, *P*=0.751), nor eggshell maculation (β±SE: −0.09±0.12, *t*_55_=-0.74, *P*=0.463).

At Day 15, the measured hormones remaining in the yolk were not related to the size of the embryos in the eggs where both were measured simultaneously. The first and second hormone principal component (Supplementary Table S6) did not explain the first embryo PC (overall size, Supplementary Table S7; hormone PC1: β±SE:0.64±0.34, *F*_1,8_=3.61, *P*=0.094, hormone PC2: β±SE:-0.09±0.53, *F*_1,8_=0.03, *P*=0.857) or the second embryo PC (embryo shape, hormone PC1: β±SE:0.17±0.25, *F*_1,8_=0.44, *P*=0.526; hormone PC2: β±SE:0.56±0.28, *F*_1,8_=4.00, *P*=0.081).

## Discussion

Eggshell maculation was not related to initial yolk steroid concentrations nor eggshell surface temperature, but eggs with higher surface temperatures on Day 9 did differ in yolk steroid hormone concentrations relative to eggs with lower surface temperatures on Day 9.

Because egg temperature varied with incubator, our experiment did not allow us to distinguish between effects of light exposure or temperature; future experiments should vary the two variables independently. Eggshell surface temperature and incubator assignment did not predict embryonic size but eggshell maculation did, as eggs with the greatest maculation produced embryos with larger bills and smaller wings on Day 15 than eggs with lower maculation.

However, both the effects of surface temperature on Day 9 hormone variation and the effects of maculation on Day 15 embryo size explained only a small proportion of the overall variation in steroid hormone concentrations or embryo size. Overall, our results provide novel evidence that eggshell maculation predicts aspects of embryonic development in birds despite maculation not being reflective of initial (Day 0) yolk steroid content in these quail, and that environmental variation may influence steroid declines without measurable effects on embryonic growth.

Of all the steroids detected in the egg, progestogens had the highest concentrations, including pregnanedione and progesterone which have previously been detected in high concentrations in other avian eggs (Merrill et al. 2019). Despite their prevalence, egg yolk progestogens have received less research attention than other steroid hormones like androgens (Groothuis et al. 2019). Progestogen injections prior to incubation can increase embryonic size in chickens early in development, although whether this has a lasting impact on hatchlings is unexplored (Paitz and Cagney 2019). Average concentrations of most of the detected yolk progestogens fell by at least 50% from Day 0 to Day 9 of incubation, with the exception of pregnanolone and progesterone. Although the progesterone concentrations varied enough among egg yolks 9 days into incubation that any differences in average concentration should be viewed with caution, it is interesting that the eggs with the warmest surface temperatures had higher concentrations of yolk progesterone remaining at Day 9, suggesting that temperature variation could be altering embryonic metabolism or uptake of this steroid. By Day 15 the concentrations of progestogens remaining in the yolk had greatly declined, although pregnanedione, pregnanolone, pregnenolone, were all present still detectable in each egg, and progesterone was still detectable in most eggs. This suggests that by ∼2 days before hatch most of the progestogens available to the embryo had been taken up and/or metabolized into a conjugate that we did not measure (Paitz and Cagney 2019), yet some were still remaining for further absorption and/or metabolism.

In addition to the progestogens, we detected historically better-known egg yolk androgens like testosterone, DHEA, and androstenedione (Groothuis et al. 2005, Hegyi and Schwabl 2010). Androgens such as testosterone and androstenedione have previously been implicated in variation in development in quail (Okuliarová et al. 2007), commercial game birds like chickens (Treidel et al. 2013), and wild songbird hatchlings like Red-winged Blackbirds (*Agelaius phoeniceus*) (Lipar and Ketterson 2000), although the direction and strength of the effects of these hormones is inconsistent (von Engelhart and Groothuis 2011). Average yolk concentrations of testosterone, DHEA, and androstenedione declined from Day 0 to Day 9 of incubation, with some Day 9 eggs no longer having any detectable DHEA or androstenedione. Like progesterone, average androstenedione and testosterone concentrations were higher in the eggs with the warmest surface temperatures on Day 9, suggesting that temperature could be altering embryonic uptake and metabolism of these hormones as well. If the variation in surface temperature measured in this study can influence hormone concentrations but not embryo size, exposure to more contrasting temperatures could induce greater differences in hormone metabolism trajectories, with implications for embryonic development. By Day 15, most eggs no longer had any detectable androgens; no eggs in the study had DHEA or testosterone on Day 15, and only one had detectable androstenedione, suggesting by ∼2 days before hatch embryos are not receiving androgens from the yolk.

We did not detect measurable levels of glucocorticoids in these quail egg yolks at any point in development (Days 0, 9, or 15) with the exception of four eggs that had low concentrations of 5β-tetrahydrocortisol on Day 9. This is surprising given the presence of glucocorticoids in past studies on commercial poultry including Japanese Quail (Hayward et al. 2006) and in avian egg yolks in several wild species (Pitk et al. 2012), including those assayed with the same hormone extraction techniques used here (Merrill et al. 2019). Because corticosterone can impact morphological development in birds (Love et al. 2005, Hayward et al. 2006) and avian corticosterone response can be impacted by pre-hatch temperature (Iqbal et al. 1990), glucocorticoids remain an intriguing mechanistic link between pre-hatch environmental variation and embryonic development. If the eggs used in this experiment did not have detectable maternally invested glucocorticoids that could link environmental conditions to embryonic development, perhaps this is one reason why we did not see effects of light treatment and temperature on embryonic growth.

Regardless of eggshell maculation or surface temperature, yolk steroid hormone concentrations consistently fell from initial concentrations to Day 15 of development (Figure 2). We did not measure any increase in yolk steroids indicating embryos’ endogenous production of hormones during development, with the exception of the 5β-tetrahydrocortisol that was only detected on Day 9. This could indicate that endogenous progestogen and androgen hormone production is not reflected in yolk levels and needs to be measured in embryonic tissue instead. The hormone measurements on Day 9 and Day 15 only reflect the concentrations in the eggs that survived to Day 9 and Day 15, as the yolks in eggs that were inviable or contained deceased embryos had started to rot. If hormone concentrations were implicated in the viability of these embryos, these data did not adequately capture the hormone concentrations that led to variation in survival. On Day 15, we were able to relate the surviving embryos’ size to the remaining detectable concentrations of steroids in the yolk, but there was no significant relationship, suggesting that there was no effect of the hormone uptake or metabolism on embryo size.

However, it is possible that the hormone variation may have implications for other aspects of embryonic or hatchling development, such as altering embryonic brain development or other organizational effects (von Engelhardt and Groothuis 2011, Groothuis et al. 2019).

Despite exposure to variation in yolk steroid hormones and hormone metabolism/uptake, surface temperatures, eggshell maculation, and light regimes, we found that Japanese Quail embryos grew to the approximately the same pre-hatch size. The morphological growth of these embryos appears to be robust to this variation in developmental environment in the first 15 days of their ∼17 day incubation period, but it is possible that the embryos could have experienced differences in development that we did not measure, such as changes in their circulating steroid hormones or cognitive development. The progestogen and androgen concentration differences seen in yolk hormones on Day 9 suggest that eggshell surface temperature could alter steroid hormone uptake and metabolism, but further research is needed to elucidate the mechanism behind this effect and its implications for hatched chicks. Nevertheless, these data provide more evidence of the presence of maternally derived yolk steroid hormones (including understudied steroids), the declines in yolk steroid concentrations with development, and the possibility that environmental inputs could alter hormone uptake and/or metabolism to explain previously established relationships between environmental variation and embryonic development.

## Statements and Declarations

### Ethical considerations

All animal handling was done in accordance with the National Research Council’s Guide for the Care and Use of Laboratory Animals and followed policies of IACUC the University of Illinois at Urbana-Champaign.

### Consent to participate

Not applicable.

### Consent for publication

Not applicable.

### Data availability

Data and script are currently available on FigShare (doi: 10.6084/m9.figshare.23590272).

## Supplementary Materials

**Table S1.** Effect of assigned incubator (24 hour light exposure or 12 hours on/off) and eggshell maculation percentage on quail yolk hormone concentrations (ng/g) of each hormone on Day 0, before the eggs were treated with lighted incubation.

| Hormone | Assigned incubator |  |  | Eggshell maculation percentage |  |  |
| --- | --- | --- | --- | --- | --- | --- |
| | $\beta \pm SE$ | $F_{1,4}$ | $P$ | $\beta \pm SE$ | $F_{1,4}$ | $P$ |
| Pregnanolone | 23.40±26.69 | 0.77 | 0.430 | 0.36±0.54 | 0.43 | 0.549 |
| Pregnanedione | 523.3±203.3 | 6.63 | 0.062 | -0.32±6.51 | <0.01 | 0.963 |
| 17α-hydroxypregnenolone | 2.70±1.89 | 2.04 | 0.226 | 0.03±0.04 | 0.51 | 0.515 |
| Pregnenolone | 78.60±38.63 | 4.14 | 0.112 | 0.04±1.08 | <0.01 | 0.969 |
| Progesterone | 27.65±20.54 | 1.81 | 0.250 | 0.35±0.45 | 0.58 | 0.488 |
| Etiocholanolone | -1.14±5.17 | 0.05 | 0.836 | 0.15±0.07 | 5.15 | 0.086 |
| DHEA | 0.12±0.66 | 0.03 | 0.870 | 0.02±0.01 | 2.92 | 0.163 |
| Testosterone | 1.06±1.06 | 0.99 | 0.375 | 0.03±0.02 | 3.47 | 0.136 |
| Androstenedione | 3.85±3.18 | 1.47 | 0.292 | 0.02±0.07 | 0.12 | 0.748 |

**Table S2.** Linear discriminant (LD) values for the first two linear discriminant analyses of the effect of eggshell surface temperature on quail yolk hormone concentrations (ng/g), with proportion of trace values indicating the amount of variation captured by each analysis and individual values for each hormone in the analysis.

|  | **LD1** | **LD2** |
| --- | --- | --- |
| *Proportion of Trace:* | *0.999* | *0.0003* |
| **Hormones:** |  |  |
| 17α-hydroxypregnenolone | -27.316 | 3.445 |
| 5β-tetrahydrocortisol | 131.799 | -3.020 |
| Androstenedione (A4) | -1.747 | 4.143 |
| Dehydroepiandrosterone (DHEA) | 147.675 | 1.902 |
| Etiocholanolone | 54.733 | 16.186 |
| Pregnanedione | 116.368 | -42.202 |
| Pregnanolone | -87.643 | -0.932 |
| Pregnenolone | 189.750 | 17.080 |
| Progesterone | -91.655 | 16.142 |
| Testosterone | 69.873 | -26.521 |

**Table S3.** Range, mean, and standard deviation of egg yolk hormone concentrations (ng/g) in viable quail eggs incubated for 9 days, separated by average eggshell surface temperature (less than and greater than 31.3 °C, the average temperature measured in the study).

|  | **Surface < 31.3 °C** | **Surface > 31.3°C** |
| --- | --- | --- |
|  | N=7 | N=3 |
| **Androstenedione (A4)** |  |  |
| range | 0–2.94 | 0–4.82 |
| mean ± SD | 0.42 ± 1.11 | 1.61 ± 2.78 |
| **Dehydroepiandrosterone (DHEA)** |  |  |
| range | 0–1.01 | 0.56–0.63 |
| mean ± SD | 0.54 ± 0.38 | 0.59 ± 0.04 |
| **Etiocholanolone** |  |  |
| range | 0.80–2.88 | 1.03–2.16 |
| mean ± SD | 1.67 ± 0.66 | 1.75 ± 0.62 |
| **Testosterone (T)** |  |  |
| range | 0.16–0.85 | 0.61–2.78 |
| mean ± SD | 0.41 ± 0.27 | 1.55 ± 1.11 |
| **17α-hydroxypregnenolone** |  |  |
| range | 0–4.66 | 0–2.68 |
| mean ± SD | 1.86 ± 2.33 | 0.89 ± 1.55 |
| **Pregnanedione** |  |  |
| range | 296–976 | 446–736 |
| mean ± SD | 571.43 ± 231.23 | 578.00 ± 146.74 |
| **Pregnanolone** |  |  |
| range | 30.0–88.4 | 52.6–124.4 |
| mean ± SD | 61.3 ± 20.0 | 79.20 ± 39.35 |
| **Pregnenolone (P5)** |  |  |
| range | 101.4–183.0 | 97.8–128.8 |
| mean ± SD | 127.23 ± 26.01 | 108.33 ± 17.73 |
| **Progesterone (P4)** |  |  |
| range | 7.64–38.20 | 13.52–37.40 |
| mean ± SD | 18.37 ± 12.01 | 29.04 ± 13.45 |
| **5β-Tetrahydrocortisol** |  |  |
| range | 0–2.94 | 0–0.3 |
| mean ± SD | 0.87 ± 1.35 | 0.10 ± 0.17 |

**Table S4.** Linear discriminant (LD) values for the first three linear discriminant analyses of the effect of incubator (exposed to either 24 hours of light or 12 hours on/off) on quail yolk hormone concentrations (ng/g), with proportion of trace values indicating the amount of variation captured by each of the three analyses and individual values for each hormone in the analysis. Bold values indicate the largest LD2 and LD3 values.

|  | **LD1** | **LD2** | **LD3** |
| --- | --- | --- | --- |
| *Proportion of Trace:* | *0.706* | *0.185* | *0.109* |
| **Hormones:** |  |  |  |
| 17α-hydroxypregnenolone | 2.839 | 4.321 | 4.986 |
| 5β-tetrahydrocortisol | -1.909 | 1.183 | -3.350 |
| Androstenedione (A4) | -9.106 | 4.628 | **-13.616** |
| Dehydroepiandrosterone (DHEA) | -8.782 | **11.074** | **-20.154** |
| Etiocholanolone | 5.043 | **-9.082** | 8.701 |
| Pregnanedione | 4.201 | **-21.402** | -6.309 |
| Pregnanolone | -0.453 | 8.291 | 6.945 |
| Pregnenolone | -17.472 | **12.025** | 6.332 |
| Progesterone | 9.772 | -0.642 | **9.022** |
| Testosterone | -0.390 | -6.814 | 4.051 |

**Table S5.**
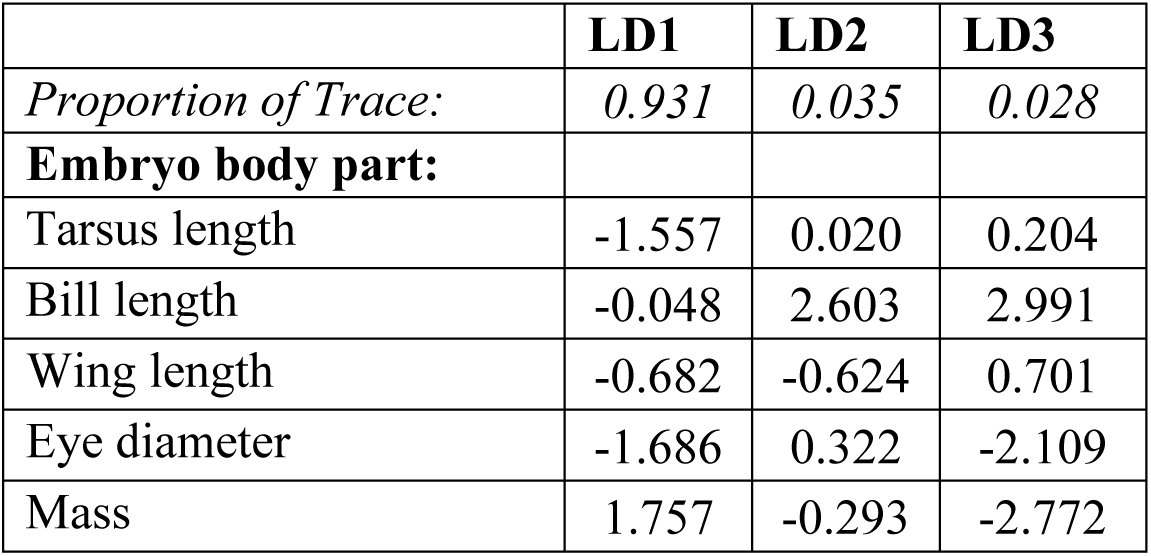
Linear discriminant (LD) values for the first three linear discriminant analyses of the effect of incubation day (9 or 15) and eggshell maculation on embryo size, with proportion of trace values indicating the amount of variation captured by each analysis and individual values for each body part in the analysis.

**Table S6.**
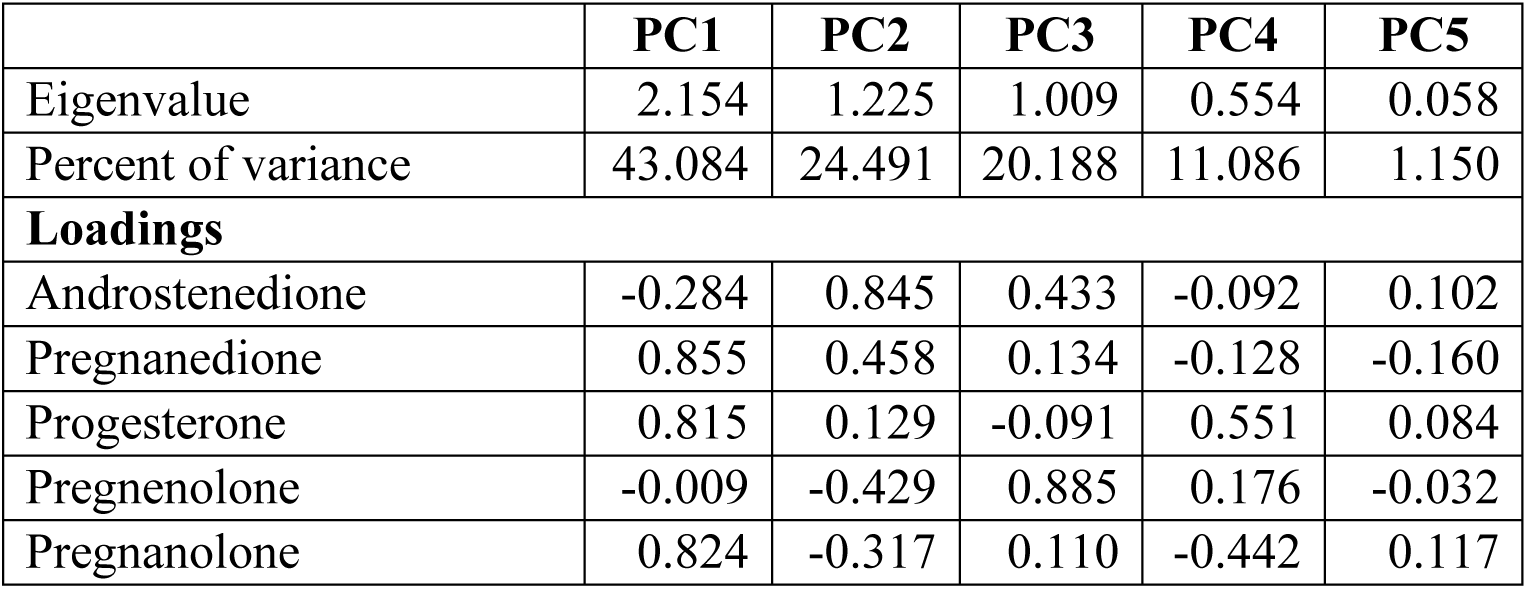
Eigenvalues, percent variance, and loadings for five principal components of hormone concentrations (ng/g) in the remaining yolk of Day 15 incubated quail eggs with viable embryos.

|  | **PC1** | **PC2** | **PC3** | **PC4** | **PC5** |
| --- | --- | --- | --- | --- | --- |
| Eigenvalue | 2.154 | 1.225 | 1.009 | 0.554 | 0.058 |
| Percent of variance | 43.084 | 24.491 | 20.188 | 11.086 | 1.150 |
| **Loadings** |  |  |  |  |  |
| Androstenedione | -0.284 | 0.845 | 0.433 | -0.092 | 0.102 |
| Pregnanedione | 0.855 | 0.458 | 0.134 | -0.128 | -0.160 |
| Progesterone | 0.815 | 0.129 | -0.091 | 0.551 | 0.084 |
| Pregnenolone | -0.009 | -0.429 | 0.885 | 0.176 | -0.032 |
| Pregnanolone | 0.824 | -0.317 | 0.110 | -0.442 | 0.117 |

**Table S7.** Eigenvalues, percent variance, and loadings for five principal components of embryo size for Day 15 viable quail embryos.

|  | **PC1** | **PC2** | **PC3** | **PC4** | **PC5** |
| --- | --- | --- | --- | --- | --- |
| Eigenvalues | 2.815 | 1.153 | 0.689 | 0.254 | 0.089 |
| Percent of variance | 56.307 | 23.059 | 13.771 | 5.083 | 1.781 |
| **Loadings** |  |  |  |  |  |
| Tarsus length (mm) | 0.427 | 0.881 | 0.138 | 0.054 | 0.141 |
| Bill length (mm) | 0.902 | -0.037 | 0.094 | -0.419 | -0.024 |
| Wing Length (mm) | 0.747 | -0.325 | -0.556 | 0.081 | 0.145 |
| Eye Diameter (mm) | 0.635 | -0.473 | 0.583 | 0.169 | 0.071 |
| Mass (g) | 0.927 | 0.216 | -0.107 | 0.201 | -0.206 |

**Figure S1.**
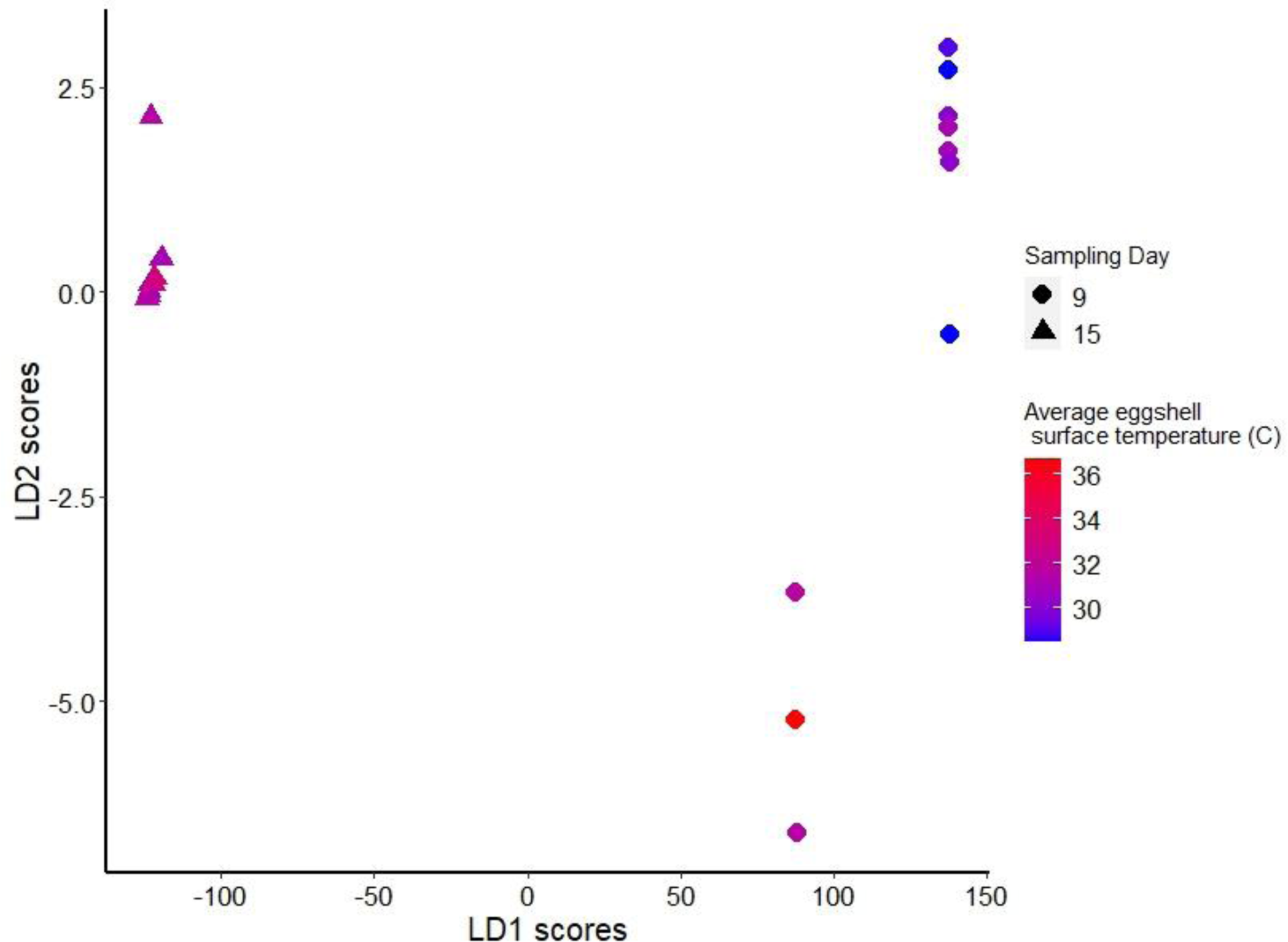
Linear discriminant (LD) scores for LD 1 and 2 illustrate that the yolk steroid hormones detected vary among viable quail eggs with different surface temperatures (⁰C). Temperatures of Day 15 eggs were averaged over measurements taken on Days 9, 12, and 15.

**Figure S2.**
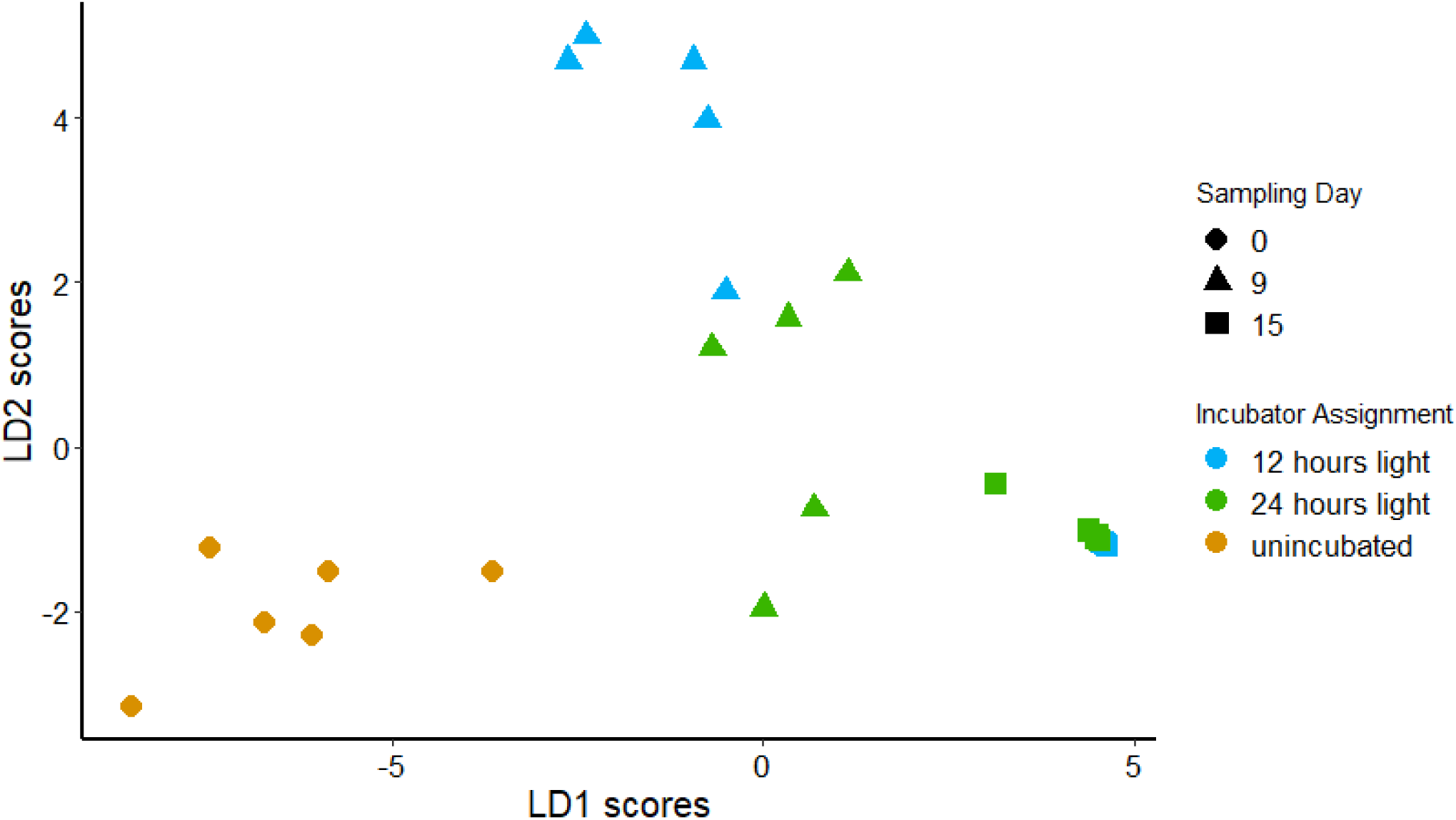
Linear discriminant (LD) scores for LD 1 and 2 illustrate that the yolk steroid hormones detected vary among viable quail eggs assigned to the two incubator treatments on Day 9 of incubation.

**Figure S3.**
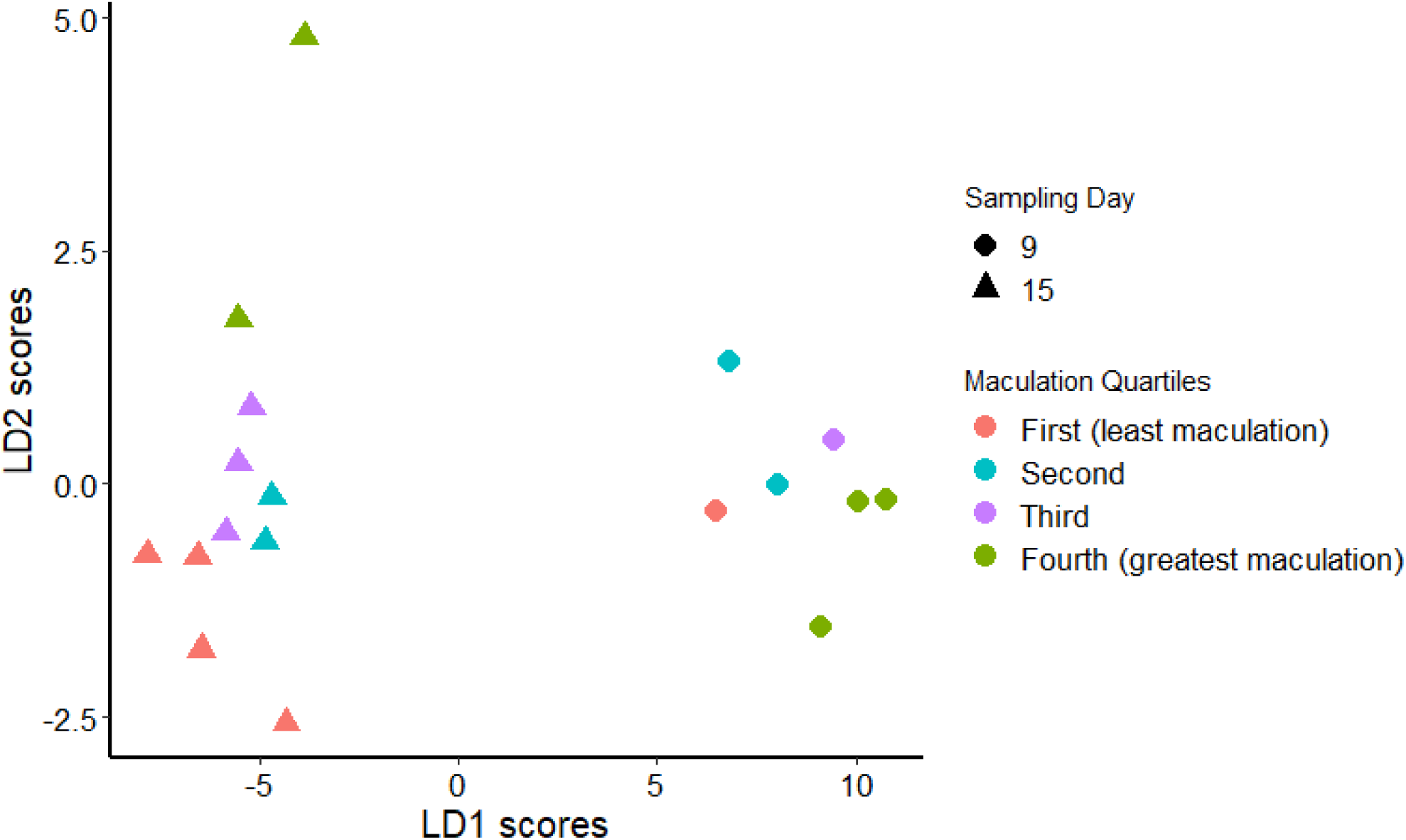
Linear discriminant (LD) scores for LD 1 and 2 illustrate that the significant effect of eggshell maculation on embryonic size measurements is driven by two eggs on Day 15 in the highest maculation quartile (most heavily spotted eggs).

**Figure S4.**
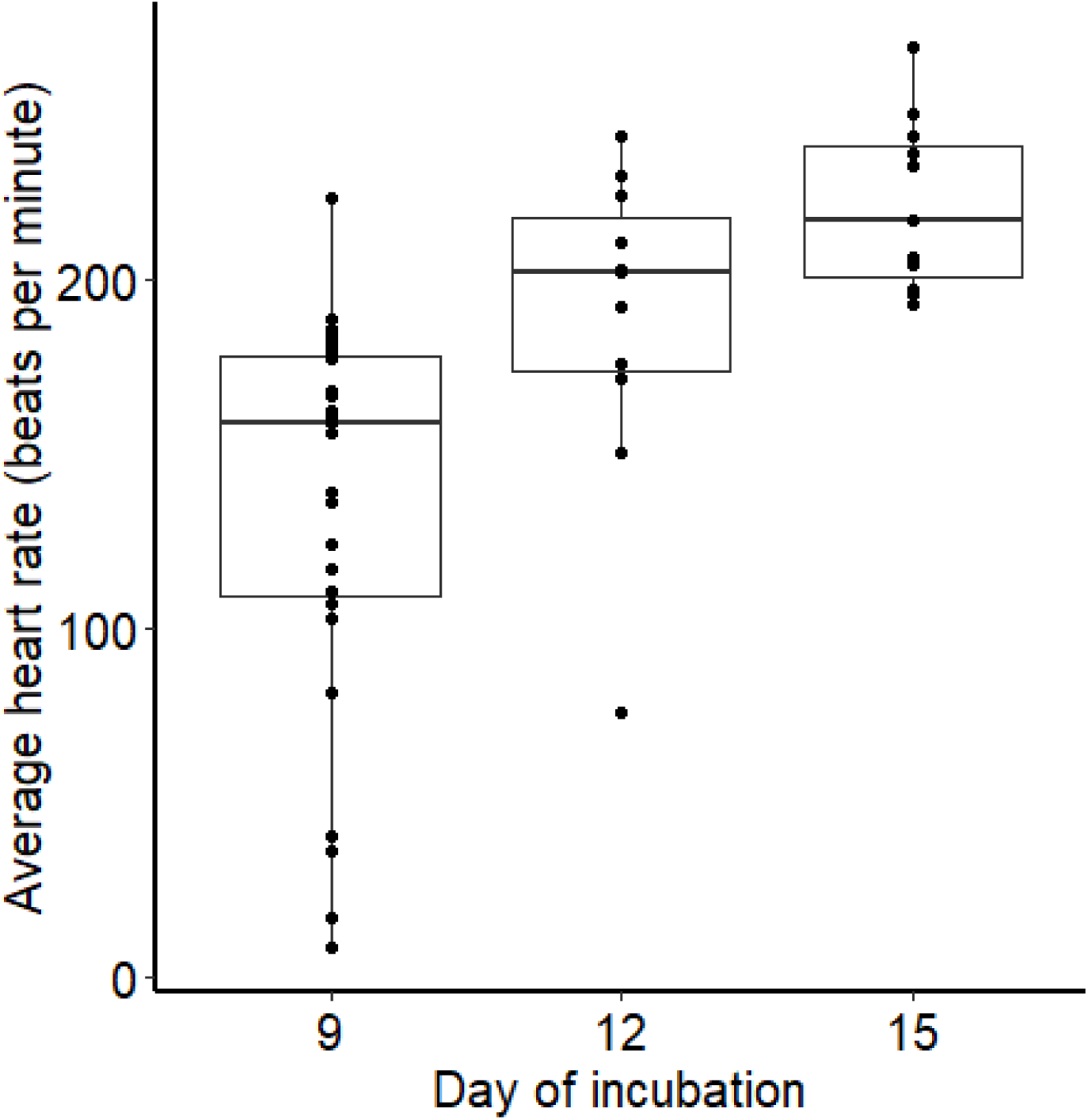
Average quail embryonic heart rate (beats per minute, sampled over five minutes) on days 9, 12, and 15 of incubation. The boxplots depict the 10^th^, 25^th^, 50^th^, 75^th^, and 90^th^ percentiles, with all data points shown as black dots for each boxplot.

